# DNA Hanger: Surface Free Single-Molecule Blotting Platform

**DOI:** 10.1101/2024.10.23.619861

**Authors:** Jincheol Seol, Byungju Kim, Eui-Sang Yu, Cherlhyun Jeong, Jong-Bong Lee

**Affiliations:** School of Interdisciplinary Bioscience & Bioengineering, Pohang University of Science & Technology (POSTECH), Pohang, Korea 37673; Department of Physics, POSTECH, Pohang, Korea 37673; Materials and Components Research Division, Electronics and Telecommunications Research Institute (ETRI), Daejeon, Korea 34129; Chemical and Biological Integrative Research Center, Korea Institute Science and Technology (KIST), Seoul, Korea 02792; Division of Bio-Medical Science & Technology, University of Science and Technology (UST), Seoul, Korea 02792

**Author notes:** To whom correspondence should be addressed. Email: C.J. or J.B.L. The first two authors should be regarded as Joint First Authors.

## Abstract

Immunoblotting typically requires protein immobilization onto a membrane surface for quantification. Particularly, in real-time single-molecule blotting, stringent surface passivation is essential to prevent false-positive signals from nonspecific binding. In this study, we introduce a novel surface-free single-molecule blotting platform, termed the DNA Hanger. This platform employs a 3D structure of a quartz slide. Modified *λ*-phage DNA molecules are randomly biotinylated along their bases and suspended between 4 μm-high thin barriers, positioned away from the quartz slide surface. A light sheet, produced within the 3D structure, illuminates fluorophore-conjugated proteins bound to the biotinylated DNA, enabling single-molecule detection. The DNA Hanger assay significantly reduces nonspecific binding and enhances sensitivity to sub-picomolar concentrations, suggesting that this platform provides a novel, surface-condition-independent, and real-time approach in a wide range of single-molecule blotting.

## INTRODUCTION

Real-time single-molecule blotting is an advanced variation of traditional blotting techniques, which have long been indispensable tools for detecting and analyzing biomolecules(1). Unlike traditional blotting(2), this cutting-edge approach offers a unique opportunity to study the dynamics and interactions of individual molecules within complex molecular assemblies, enabling a comprehensive understanding of their functional roles and regulatory mechanisms(3,4). The real-time aspect comes from the ability to track individual molecular events using high-sensitivity fluorescence microscopy and single-molecule detection technologies(5).

In this technique, individual biomolecules are immobilized onto a surface, typically a membrane or specialized substrate. Ideally, target molecules should bind to specific sites on the surface, but, in practice, they also adhere to the surface nonspecifically. Such nonspecific binding can distort the signal, leading to erroneous interpretations of molecular behaviors and interactions. Although there are several strategies to reduce nonspecific binding to the surface using surface blocking with agents like Polyethylene glycol and lipid bilayer or altering the charge or hydrophobicity of the surface(6-9), reducing nonspecific binding in real-time single-molecule blotting remains a critical challenge that can interfere with the accuracy and reliability of the assay(10).

Recently, we have developed a single-molecule fluorescence imaging platform to minimize interference from nonspecifically surface-bound proteins(11). This platform is designed to monitor the dynamics of individual fluorescently labeled proteins on DNA molecules away from the surface. The platform features thin quartz barriers a few micrometers in height. The λ-phage DNA molecules (48.5 kilo-base pairs) are biotinylated randomly to their bases, where target proteins and/or antibodies are immobilized. The modified DNA molecules were stretched and suspended across two adjacent barriers using laminar flow, which aligned the DNA along these ridges. A thin light sheet produced near the ridges due to interference from excitation beams refracted by the barriers enables us to visualize individual fluorophores with high precision(11). Using the platform, we show that extremely low nonspecific bindings allow for measuring the quantity of proteins of interest simply by counting the number of its fluorophore-conjugated antibody or fluorophore-labeled target proteins with high specificity and sensitivity.

## MATERIALS AND METHODS

### 3D Patterned Quartz Slide

The fabrication of 3D-etched quartz slides was performed with slight modifications to a previously established protocol(11). Briefly, quartz wafers were thoroughly cleaned using a piranha solution (a 3:1 mixture of H_2_SO_4_ to H_2_O_2_) at 130 °C for 10 minutes. Following this, an adhesion promoter (hexamethyldisilane, J.T.Baker) and a photoresist (TDMR-AR87, Tokyo Ohka Kogyo) was applied to the surface through spin coating at 3,000 rpm and 4,500 rpm, respectively, then baked at 88°C for 60 seconds. A stripe pattern was created on the quartz surface using soft contact mode with a photomask aligned by an MA-6 mask aligner and rinsing in a developer (AZ 300 MIF, Merck) for 60 seconds. The patterned surface underwent a hard bake at 113°C for 90 seconds to develop the design. The final step involved etching the quartz surface by immersing it in a 6:1 diluted HF solution (Buffered oxide etch Samchun) at room temperature without stirring. This fabrication process produced quartz slides with the following dimensions: barrier height of 4 μm and barrier intervals of 13.1 μm.

### Quartz slide surface cleaning and passivation

The quartz slides were subject to a rigorous washing to ensure complete removal of impurities. The cleaning process began with incubation in a staining jar containing 10% Alconox for 20 minutes, followed by sequential washes in ddH_2_O for 5 minutes, acetone for 15 minutes, KOH for 20 minutes, piranha solution for 20 minutes, and methanol for 10 minutes. Sonication was employed during these procedures to enhance impurity removal. After the washing steps, the slides were briefly exposed to a flame using a torch for 30 seconds to ensure thorough surface cleaning.

For PEG surface passivation, the cleaned quartz slides were incubated in a 2% (v/v) solution of (3-aminopropyl)triethoxysilane (a3648, Sigma-Aldrich) for 20 minutes. A mixture of PEG-succinimidyl valerate (PEG-SVA, MW 5000, Laysan Bio) and biotin-PEG-succinimidyl valerate (MW 5000, Laysan Bio) in a 0.1 M sodium bicarbonate solution (pH 8.3) was prepared at a 40:1 mass ratio. The PEG solution was carefully applied to the surface of the quartz slides and incubated overnight. After incubation, the PEG-passivated quartz slides were thoroughly rinsed with deionized water, and any residual water was removed from the surface using high-pressure nitrogen gas.

### Anti-digoxygenin antibody stamping with PDMS

A mixture of PDMS base and curing agent (7:1, SYLGARD® 184 SILICONE ELASTOMER KIT, Dow Corning) was poured onto a flat plate. The mixture was degassed in vacuum desiccator (∼0.2 atm) until all air bubbles were eliminated. The plate was then baked in an oven at 80°C for 1 hour to cure the PDMS. After the curing process, the PDMS was cut into pieces matching the dimensions of the pattern on the quartz slide, with a size of 11mm × 4mm.

To prepare the stamping side on the PDMS pieces, 10 μl of anti-digoxigenin solution (200 μg/ml, diluted in PBS; 11333089001, Roche) was placed and spread on the PDMS surface inside a moisture box. After a 30-minute incubation, the solution on the PDMS surface was gently removed by nitrogen gas. The prepared PDMS stamp was then carefully applied onto the patterned surface of the quartz slide. Finally, the quartz slide and a cover glass were assembled into a flow chamber using double-sided tape.

### Biotinylated-long DNA

The DNA substrate was prepared using bacteriophage λ-phage DNA (48.5 kb, New England Biolabs) through ligation with 14 nt-length 3’ digoxygenin-labeled oligonucleotides, using T4 ligase (NEB) at room temperature overnight (Table S1). After ligation, excessive oligonucleotides were removed by dialysis (Spectra-Por Float-A-Lyzer®, G2) or ultrafiltration (10 kDa Amicon, Millipore).

Subsequently, biotin was conjugated to the λ-phage DNA using either photobiotin (PHOTOPROBE®, Vector) or psoralen-biotin (HOOK™-Psoralen-PEO-Biotin, G-biosciences). The biotin reagent solution was added to the purified λ-phage DNA solution in a 50:1 volume ratio for photobiotin and a 100:1 for psoralen-biotin. The solutions were gently mixed by inverting the tubes, and the tubes were placed 2 cm below a 365 nm hand-held mercury lamp (8 W, Sanyo) for 1 hour to facilitate the photo-coupling reaction.

In this imaging condition, purification of the DNA after the photo-coupling reaction was not a mandatory step. Additionally, we explored the use of long-range PCR (LA Taq DNA polymerase, Takara) with biotinylated dUTP (Biotin-11-dUTP Solution, Thermo Scientific™) as an alternative method for fabricating the DNA substrate. However, this method resulted in low yield due to the lengthy DNA template used in the reaction.

### mNeonGreen-PABP

The pQE31 vector expression system was used to construct plasmids for the recombinant protein expression of full-length human PABPC1(PABP) (12). The mNeonGreen (mNG) sequence was amplified by PCR from a plasmid (addgene, #98877), and the PCR product was inserted at the N-terminus of the pQE31 expression vector. Additionally, a 3xFLAG tag was subcloned into the C-terminus of the construct. The PABP gene, obtained through PCR amplication, was then inserted between mNeonGreen and the 3xFLAG sequence in the plasmid.

Recombinant proteins were overexpressed in NEBExpress® Iq Competent *E. coli* (C3037I, NEB) cultured in LB medium. Protein expression was induced by adding 0.4 mM IPTG, and the culture was incubated for 4 hours at 37°C. After cell harvesting, the recombinant proteins were purified following the manufacturer’s instructions for the NI-NTA Fast Start Kit (30600, QIAGEN) and Anti-DYKDDDDK Affinity Resin (101274-MM13-RN, Sino Biological). All steps of the protein expression and purification procedures were carried out on ice to preserve protein stability. The concentration of purified recombinant protein was measured using a nanophotometer (IMPLEN). Proteins were aliquoted and stored at -80°C.

### RNA/DNA partial duplex with poly(A) tail

To construct the partial duplex, single-stranded polyadenine RNA (114 nt) was synthesized using an *in vitro* transcription assay. Initially, the pBluescript SK(-) vector was digested at two restriction sites and ligated with oligonucleotides to form the polyadenine DNA construct template (Table S1). After ligating the oligonucleotide to the vector, the remaining gap on the upper strand was filled with T4 DNA polymerase (M0203S, NEB) according to the manufacturer’s protocol. The resulting template DNA was iterated twice to reach a poly(A) length of 114. The final RNA template was synthesized using the HiScribe™ T7 Quick High Yield RNA Synthesis Kit (E2050S, NEB), following the manufacturer’s instructions. The construct used was 5’-ctcgaggtcgacggtatcgataagcttgaagac(a)_25_-3’ and the complementary DNA oligonucleotide was modified with Cy5 at the 5’ end and biotinylation at the 3’ end (IDT, Coralville, USA).

The partial duplex was prepared by annealing the DNA oligo with the RNA transcript at a 1:9 molar ratio in annealing buffer (1 mM MgCl_2_, 10 mM Tris-HCl, 100 mM NaCl, pH 7.4) to a final concentration of 100 nM. The mixture was incubated at 85°C for 10 minutes and then allowed to cool gradually to room temperature. The resulting partial duplex (Cy5-DNA/RNA duplex) was stored at 4°C for further use.

### Fluorophore-labeled antibodies and TNF-α

For the preparation of fluorophore-conjugated antibodies, we followed the protocol described in a previous study (13). Briefly, purified human TNF-α antibodies (MAb1 and Mab11, BioLegend) were modified for either capture or detection of TNF-α. MAb1, the capture antibody (cAb), was site-specifically biotinylated using the SiteClick™ Antibody Labeling Kit (Thermo Fisher Scientific) according to the manufacturer’s instructions. MAb11, the detection antibody (dAb), was buffer-exchanged with PBS, and both MAb1 and MAb11 were separately mixed with 5 μl of Alexa Fluor™ 647 NHS Ester or Alexa Fluor™ 568 NHS Ester (Thermo Fisher Scientific) in DMSO (10 mg/ml) for 1 hour at room temperature, respectively. The modified antibodies were subsequently purified using PD MiniTrap G-25 (GE Healthcare) and buffer-exchanged with PBS containing 0.09% sodium azide using an Amicon® Ultra-10k Centrifugal Filter Unit. The concentration and degree of labeling of the antibodies were measured using a Nanophotometer. All antibodies were stored at 4°C. TNF-α (5 µg, R&D Systems) was reconstituted in PBS to 5 µM and stored at -80°C.

### Single-molecule imaging

The flow chamber was mounted on a custom-built prism-type total internal reflection fluorescence (TIRF) microscope (60× water-immersion objective, NA = 1.2, Olympus IX-71). Samples were illuminated based on the excitation wavelengths of their conjugated fluorophores: mNeonGreen at 488nm, Cy3 and SYTOX Orange (Invitrogen) at 532 nm, Alexa Fluor™ 594 at 561 nm, and Cy5 at 638 nm. All fluorescence signals were captured using an EMCCD camera (ImagEM C9100-13, Hamamatsu) and processed with camera link frame grabbers (AS-FBD-1XCLD-2PE4L-F, Active Silicon). Video recording was conducted at a rate of 100 ms per frame, which was the standard acquisition rate employed throughout this study. colocalization analysis was performed using the ImageJ (National Institutes of Health, Bethesda, MD) plugin, ComDet v.0.5.5.

Prior to image acquisition, DNA binding buffer supplemented with 5 mM PCA (P5630, Sigma-Aldrich), 200 nM rPCO (46852004, Oriental Yeast), and 2 mM Trolox were injected to reduce photobleaching and photoblinking during imaging (14). After each injection, the flow chamber was thoroughly washed with 250 µL of 1x DNA binding buffer, and the washing process was repeated twice.

## RESULTS

### DNA Hanger Assay

Digoxignenin-labeled oligonucleotides were introduced at both *cos* ends of the *λ*-phage DNA that was incorporated with biotin using photobiotin or psoralen-biotin (Figure 1a; Table S1; Materials and Methods). The modified DNA substrates were stretched and tethered between the tops of barriers that is 4 μm-high from the quartz surface via a digoxigenin-antidigoxigenin antibody interaction. To achieve the attachment, a solution of the modified *λ*-phage DNA molecules (20 pM, diluted in 1x TE buffer, pH 7.4) was injected into the flow channel at a flow rate of 0.07 ml/min with a syringe pump (Harvard Apparatus). After tethering, 500 μl of Cy5-streptavidin (20 ng/ml, S-888, Invitrogen) or Cy5-streptavidin (20 ng/ml, SA1011, Invitrogen) solution was injected into the flow chamber constructed with the 3D structure quartz slide and then incubated for 5 minutes to coat the DNA substrates containing biotin molecules (Figure 1b). This 3D structural organization of the DNA image substrate on the barriers is named DNA Hanger.

**Figure 1.**
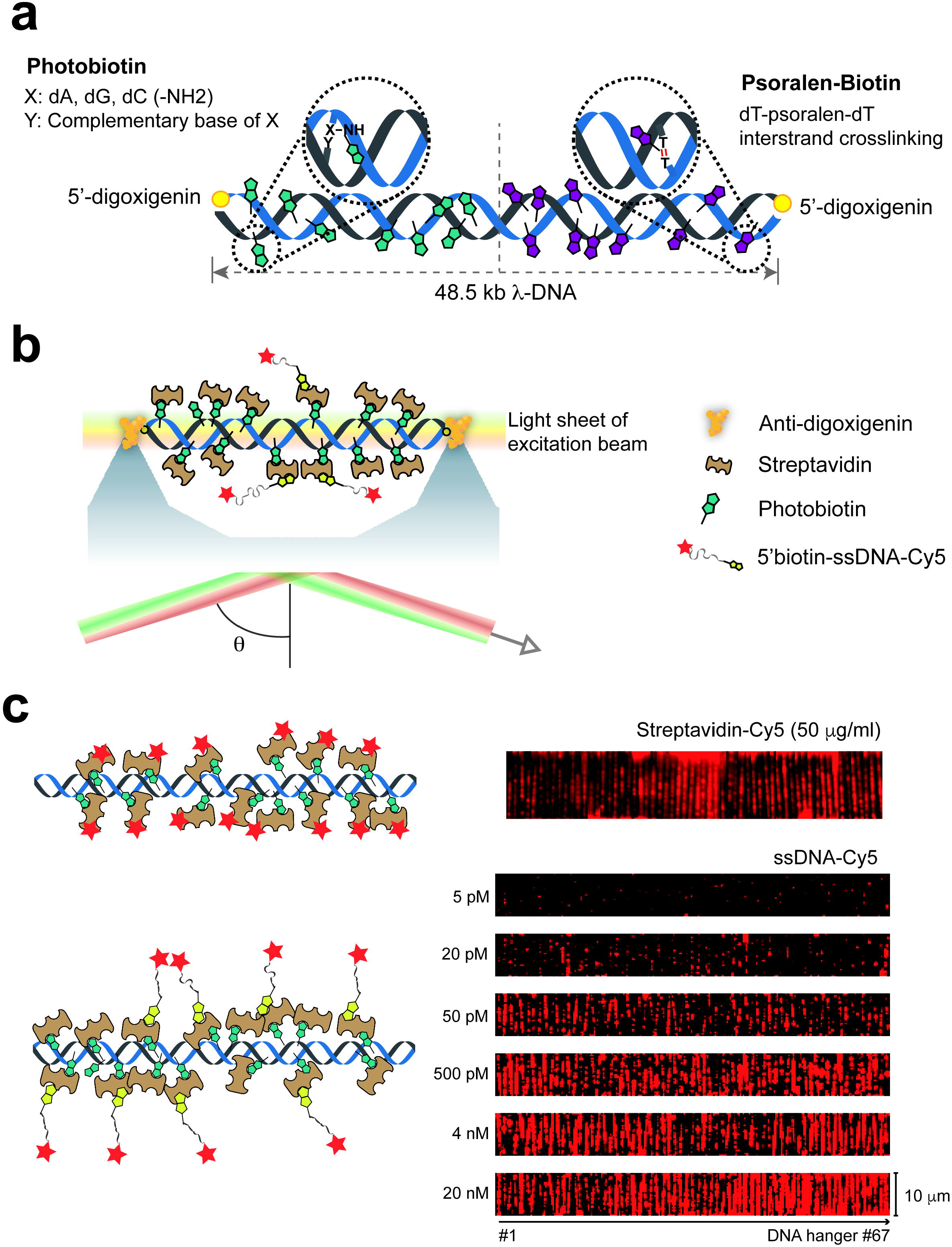
DNA Hanger Assay. **(a)** Schematic of the biotinylated DNA hanger. Biotins were generated by photo-coupling reagent (Photobiotin), which introduces biotin-group onto the bases on nucleic acids upon 405 nm irradiation. A 3’-digoxigenin oligonucleotide was ligated to 12 nt cohesive ends of the linearized λ-phage DNA. **(b)** Due to the 3D-etched quartz surface, the excitation laser forms a thin light sheet near the barrier where the biotinylated DNA is tethered via a streptavidin and anti-digoxigenin antibody interaction. Target molecules (5’-biotin-ssDNA-Cy5) are immobilized on the DNA Hanger through the interaction between streptavidin and biotin. **(c)** Schematic of DNA Hanger (upper left) and representative images with Cy5-streptavidin (upper right). Schematic of DNA Hanger imaging with unlabeled streptavidin and Cy5-labeled DNA (bottom left), along with representative images of the DNA Hanger displaying various concentrations of Cy5 labeled DNA (bottom right).

In our previous study (11), we demonstrated that the unique 3D structure of the quartz slide could generate a light sheet with a full width half maximum of about 1.1 μm from the incident beam, depending on the incident angle. This light sheet formed an image plane on top of the barriers when the excitation beam was a incident angle of *θ* ∼ 75° (Figure 1b). This configuration dramatically reduces background signals from analytes or ligands that nonspecifically bind to the bottom surface, as emitters on the bottom surface appear to be out of focus. Consequently, the signal-to-noise ratio of the probe on the DNA imaging substrate is sufficiently high to allow for the visualization of individual fluorophores.

We observed fluorescence signal of the Cy5 emitters along the DNA Hanger, confirming that biotin-streptavidin interaction was occurred on our modified λ-phage DNA. Notably, the DNA imaging substrate was saturated with Cy5-labeled streptavidin at a concentration of 50 μg/mL (Figure 1c top). This complete occupation of the DNA by streptavidin can be leveraged to control the density of ligands for subsequent binding studies.

To assess the binding capacity and unbiased nature of the DNA Hanger assay, we conducted binding experiments using Cy5-ssDNA of 100 nt (Table S1) across a concentration range from 5 pM to 20 nM, which were diluted in DNA binding buffer (pH 7.5, containing 30 mM Tris-HCl, 100 mM potassium glutamate, 5 mM MgCl2, and 0.0025% Tween-20). We observed that as the concentration of injected Cy5-labeled ssDNA increased, the DNA Hanger’s binding capacity reached saturation at approximately 4 nM ssDNA (Figure 1c). Our results demonstrate the successful development and validation of the DNA Hanger platform for precise molecular tethering and analysis in wide range of single-molecule blotting assays.

### Real-time single-molecule immunoblotting using DNA Hanger

Real-time single-molecule immunoblotting represents a significant advancement in the study of molecular interactions, but it often suffers from severe nonspecific antibody binding, which impedes precise quantitative analysis. Despite employing strigent surface passivation, this method does not always yield consistent or homogenous results across different trials in a conventional PEG-biotin quartz slide, highlighting the limitations of tranditional approaches in minimizing nonspecific binding.

To assess the usability of the DNA Hanger for single-molecule immunoblotting, we examined the binding of PABP to poly(A) RNA. An RNA substrate containing 114 nt poly(A) was annealed with DNA oligonucleotides labeled with Cy5 at the 5’ end and biotinylated at the 3’ end. The Cy5-DNA/RNA partial duplex with a poly(A)_114_ tail was diluted to 5 pM in reaction buffer (10 mM Tris-HCl, pH 8.0, 100 mM KCl. 5 mM MgCl_2_). The DNA/RNA partial duplex solution was then injected into the flow chamber. The resulting DNA/RNA partial duplex was then tethered to either the DNA Hanger or a conventional PEG-biotin quartz slide.

We introduced 100 nM mNG-PABP into the chamber, followed by washing to remove unbound mNG-PABP molecules (Figure 2a). Subsequently, 53.3nM Alexa fluor® 594-conjugated PABP antibody (AF594-PABP antibody; (10E10, Santa Cruz)) was added to the channel containing mNG-PABP bound to the poly(A)_114_ of the Cy5-DNA/RNA partial duplex in the DNA Hanger and the chamber was washed again to remove free antibodies in solution (Figure 2a).

**Figure 2.**
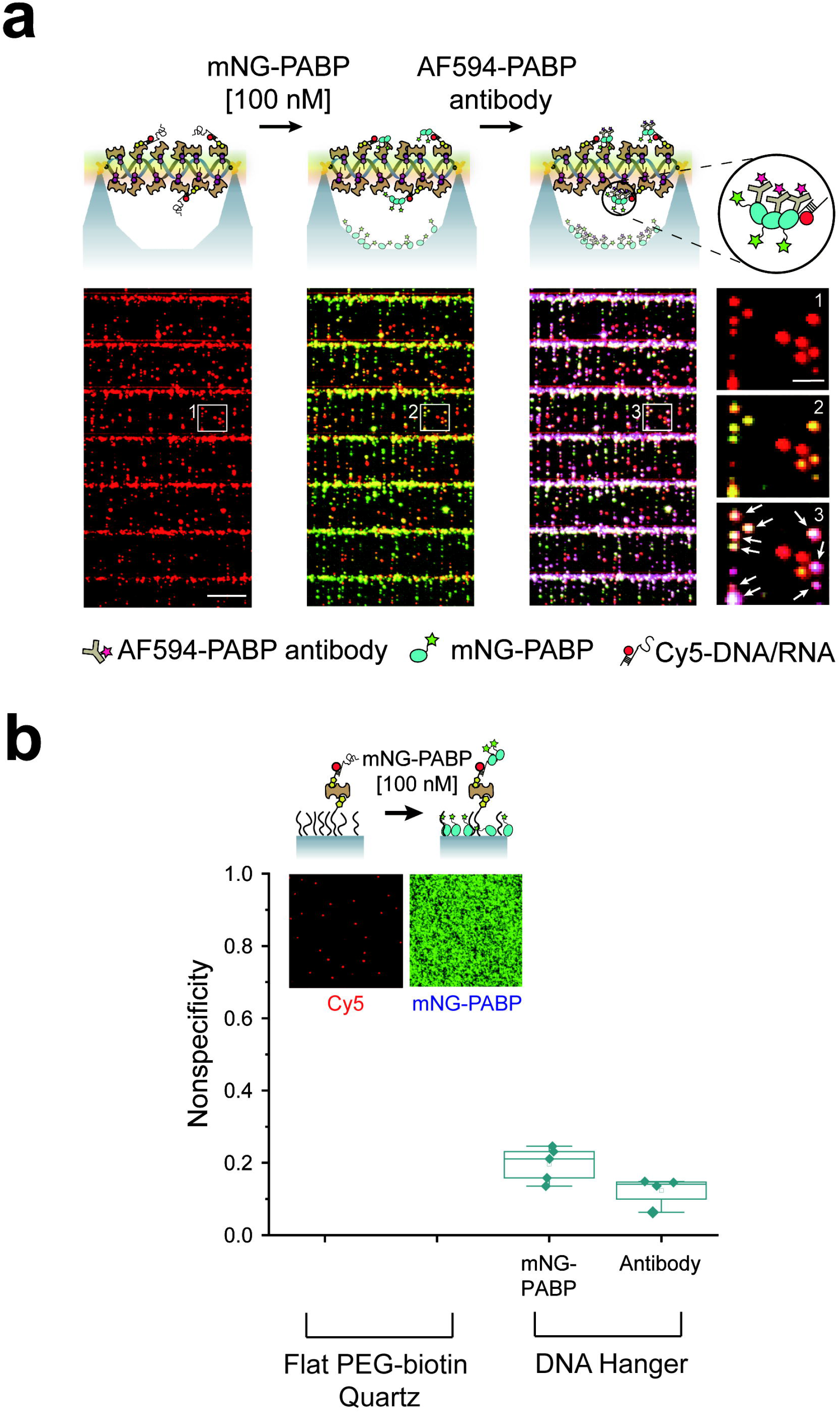
PABP-poly(A) RNA binding. **(a)** Schematic (upper) and corresponding representative fluorescence image (lower) of a Cy5-DNA/RNA partial duplex (red) tethered to the DNA Hanger through streptavidin-biotin interaction (left), mNG-PABP (green) bound to the Cy5-DNA/RNA partial duplex (middle), and AF594-conjugated PABP antibody (pink) bound to mNG-PABP (right). A magnified view of the white boxed area is shown on the far right. White arrows indicate spots where colocalization of all three colors occurs. The Cy5-DNA/RNA partial duplex containing poly(A)_114_ in this image. **(b)** Nonspecific binding ratio of mNG-PABP to AF594-conjugated PABP antibody in the DNA Hanger. Error bars indicate standard deviations (s.d.).

Using colocalization analysis on the DNA Hanger (15), we measured the nonspecific binding of mNG-PABP (0.20 ± 0.05; mean ± s.d.) and AF594-PABP antibody (0.12 ± 0.04) as shown in Figure2b. The nonspecificity represents the ratio of mNG-PABP not colocalized with the Cy5-DNA/RNA partial duplex and the ratio of AF594-PABP antibody non colocalized with mNG-PABP to the total number of mNG-PABP or AF594-PABP antibody (N = 1339 for mNG-PABP, N = 1003 for AF594-PABP antibody). Due to excessive nonspecific binding of mNG-PABP on the PEG-biotin quartz slide even after rigorous washing (Figure 2b), accurate analysis was not available. This high level of nonspecific binding hinders to perform reliable quantitative analysis on the conventional PEG-biotin surface.

The 20% and 12% nonspecific binding observed on the DNA Hanger could be attributed to photobleaching or the presence of non-fluorescent Cy5 in the DNA/RNA duplex and mNG-PABP. In a control experiments, we barely detected nonspecific binding signals of mNG-PABP or AF594-PABP antibody in the absence of the Cy5-DNA/RNA partial duplex or mNG-PABP on the DNA Hanger. These findings strongly suggest that the DNA Hanger assay offers a superior platform for real-time single-molecule immunoblotting methods.

### Single-molecule ELISA with DNA Hanger

The enzyme-linked immunosorbent assay (ELISA) is a widely used technique for detecting and quantifying specific protein biomarkers by employing antibodies linked to an enzyme, ensuring both specificity and sensitivity (16). Here, we present the DNA Hanger assay as an application of ELISA for extensively studied human tumor necrosis factor α (TNF-α).

For the DNA Hanger ELISA assay, we prepared TNF-α and its fluorophore-labeled antibodies for capture (AF647-MAb1; cAb) and detection (AF568-MAb11; dAb) as described in Methods and Materials. A 7.2 nM cAb was injected into the DNA Hanger chamber and incubated for 5 minutes to ensure full coverage. After removing any excess cAb, 3-100 pM TNF-α was infused and maintained for 5 minutes. After the removal of unbound TNF-α, 40 nM dAb was added and incubated for 10 minutes (Figure 3a).

**Figure 3.**
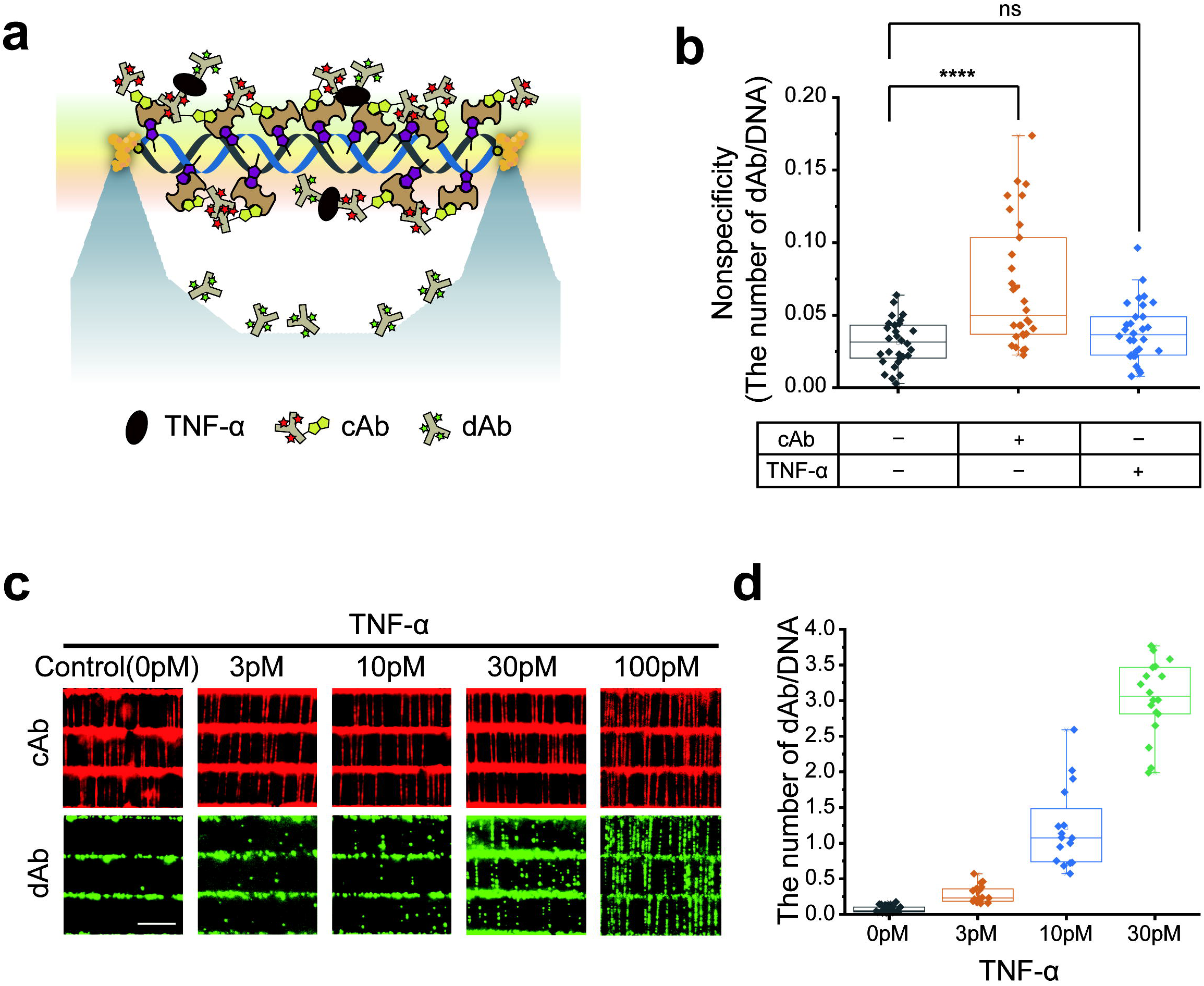
Single-molecule ELISA for TNF-α. **(a)** Schematic of TNF-α and its antibodies (cAb or dAb) attached to the DNA Hanger. The cAb, conjugated with AF647 and biotin, is immobilized on λ-phage DNA through the biotin-streptavidin interaction, and captures TNF-α, which is detected by a fluorescently labeled dAb. **(b)** The nonspecific binding of dAb or TNF-α to the DNA Hanger or cAb. Each diamond dot indicates the average of the number of dAb per DNA in a field of view. A total 10,784 DNA molecules (left), 9,954 (middle), and 6,894 (right) across 30 fields of view were analyzed, respectively. *P-*values were calculated using one-way ANOVA test (*\*\*\*\*P* < 0.00005). “ns” indicates no statistically significant difference (*P* ≥ 0.05). **(c)** Representative fluorescent images of cAb exhibiting DNA molecules and dAb binding to TNF-α across different concentrations, ranging from 0 pM to 100 pM. **(d)** The number of dAb per specific length (10 μm) of DNA was calculated at various concentrations. A total of 3,121 DNA molecules in 18 fields of view (0 pM), 3,465 in 17 fields of view (3 pM), 2,173 in 16 fields of view (10 pM), and 1,699 in 18 fields of view (30 pM) were analyzed, respectively. All data were collected from at least three independent experiments.

We assessed the degree of nonspecific bindings of dAb by quantifying the number of dAb molecules bound to the DNA Hanger (Figure 3b). The nonspecific binding of dAb in the absence of both cAb and TNF-α was negligible (0.031 ± 0.016; mean ± s.d.). When cAb was introduced without TNF-α, nonspecific binding of dAb remained extremely low (0.070 ± 0.043). The nonspecific binding is nearly identical to that without both cAB and TNF-α (0.038 ± 0.020) when TNF-α was added in the absence of cAb, which indicates that TNF-α did not bind to streptavidin coated DNA (Figure 3b). Additionally, nonspecific binding of dAb was independent of incubation time of dAb, showing 0.063 ± 0.025 after 4 hours of incubation (Supplementary Figure S1). We estimate that one dAb per 185 μm was nonspecifically bound to the DNA, taking into account the 13 μm distance between barriers and a nonspecific binding rate of 0.07 per DNA for dAb in the presence of cAb. We conclude that nonspecfic binding to the DNA Hanger was negligible.

Next, we explored the specificity of dAb binding to TNF-α in the presence of cAb at various concentrations (Figure 3c and 3d). The average number of dAb per DNA was 0.28 ± 0.13 at 3 pM TNF-α, which increased approximately 10-fold at 30 pM TNF-α (3.04 ± 0.53; Figure 3d). At concentrations higher than 30 nM, accurate counting of dAb became challenging due to the dense binding of dAb (Figure 3c). Alternatively, the fluorescence intensity of AF594-dAb offers a viable method for quantifying TNF-α. These findings demonstrate that the DNA Hanger assay is capable of detecting and quantifying target proteins at concentrations as low as subpicomolar levels in the sample.

## DISCUSSION

In this study, we developed and demonstrated the DNA Hanger assay, a novel real-time single-molecule blotting technique. This platform offers a significant advancement in both specificity and sensitivity for molecular detection and quantification, particularly in the context of single-molecule immunoassays. Compared to traditional techniques, which often suffer from high levels of nonspecific binding, the DNA Hanger assay substantially minimizes background noise, allowing for more accurate quantification of target molecules.

The most notable improvement of the DNA Hanger assay is its ability to reduce nonspecific binding. By suspending DNA molecules away from the surface on thin quartz barriers, we achieved nonspecific binding rates as low as one protein per 185 μm of DNA, even under prolonged incubation conditions. This improvement is essential in overcoming the challenges posed by nonspecific binding in traditional blotting assays, which can lead to erroneous interpretations of molecular interactions. Additionally, the DNA Hanger assay demonstrated high sensitivity. Our single-molecule ELISA results with TNF-α and its antibodies show that the platform is suitable for precise quantitative analysis with protein concentrations as low as subpicomolar levels.

Another key application of the DNA Hanger assay is its use in real-time single-molecule imaging. By leveraging the reduced nonspecific binding, we successfully quantified mNG-PABP binding to poly(A) RNA in a manner not achievable with conventional PEG-biotin quartz slides without interference from nonspecific binding. This was achieved through the photobleaching steps of poly(A) tails in two different lengths (Supplementary Figure S2).

The biotinylated long DNA, which serves as the imaging substrate in the DNA Hanger, carries a large negative charge due to its phosphate backbone, making it susceptible to nonspecific binding of positively charged molecules. We confirmed that 5 mM Mg^2+^ effectively suppressed nonspecific binding by neutralizing the charge interaction between DNA and Mg^2+^. To minimize the charge effect, other polymers can be alternative for the DNA Hanger instead of DNA. For instance, multi-walled carbon nanotubes functionalized with reactive groups, which are normally charge neutral, could be a potential candidate for immobilizing antibodies or target molecules while minimizing charge-induced nonspecific binding (17).

In conclusion, the DNA Hanger assay represents a versatile and powerful tool for single-molecule studies such as ELISA and pull-down assays. Its ability to minimize nonspecific binding and provide high sensitivity makes it an attractive platform for a broad range of applications, from basic research in molecular biology to potential clinical diagnostics. Moving forward, we anticipate that this method will open up new possibilities for exploring the detailed mechanisms of biomolecular interactions and will contribute to the development of more accurate and reliable assays for biomolecule detection.

## DATA AVAILABILITY

All data presented in this study are accessible within the main and supplementary figures.

## SUPPLEMENTARY DATA

Supplementary Data are available at NAR online.

## ACKNOWLEDGEMENTS

We thank laboratory members for helpful discussions.

