## Supplementary Information for "DNA Hanger: Surface Free Single-Molecule Blotting Platform"

**Supplementary Table 1. Oligonucleotides for DNA substrates. PAGE or HPLC-purified oligonucleotides were purchased from Bionics (http://www.bionicsro.co.kr/)**

| Bi-dig lambda DNA preparation | |
|---|---|
| **Name** | **Sequence** |
| λ-Cos_comp_dig (1) | 5'-phos/ AGG TCG CCG CCC TT/dig-3' |
| λ-Cos_comp_dig (2) | 5'-phos/ GGG CGG CGA CCT TT/dig-3' |
| **In vitro Transcription** | |
| **Name** | **Sequence** |
| Oligo_up (1) | 5'-phos/ TCG ACG GTA TCG ATA AGC TTG AAG ACA AAA A -3' |
| Oligo_up (2) | 5'- AAA AAG AGA CGC GAA TTC CTG CA -3' |
| Oligo_down_poly(T)_34 | 5'-phos/ GGA ATT CGC GTC TC $T_{(34)}$ GTC TTC AAG CTT ATC GAT ACC G -3' |
| **SSB binding experiment** | |
| **Name** | **Sequence** |
| 5'Biotin_3'Cy5_100mer_template (purchased from IDT) | /5Biosg/ GTT ACC GAT ACG ATA CGA ATA GGC ATA TCT GCA CGT TTC TCA CGA GGC GCC GCT AGA CTG ATC TGG AGC TTA ATT GCC TGC CGG AGC TAA AGA CGT TCC A /3Cy5Sp/ |

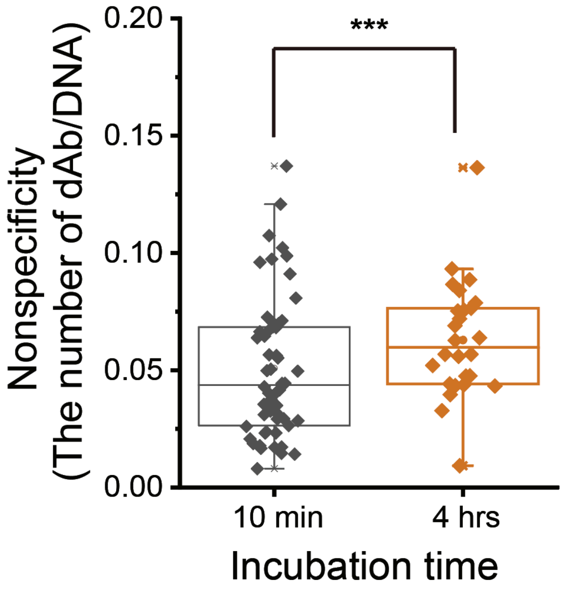


Figure S1. Nonspecific binding ratios at different incubation times. Each diamond represents the average ratio in a field of view. A total of 10,964 DNA molecules across 54 fields of view (10 min) and 2,992 DNA molecules across 26 fields of view (4 hours) were analyzed. *P*-values were calculated using a two-tailed Student's t test with unequal sample variance (****P* < $4.7x10^9$).

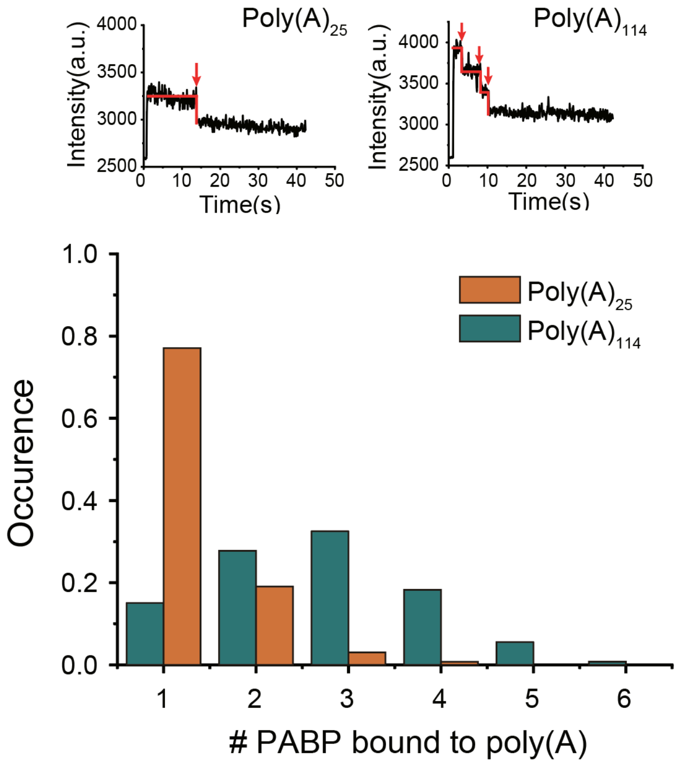


Figure S2. Upper panels: Representative intensity-time traces showing photobleaching steps of mNG-PABP bound to the poly(A) tail in the Cy5-DNA/RNA partial duplex; 1 step at poly(A)$_{25}$ or 3 steps at poly(A)$_{114}$. Lower panel: The histogram of the number of mNG-PABP bound to poly(A)$_{25}$ (orange, n = 126) or poly(A)$_{114}$ (dark turquoise, n = 131).
